# Specific patterns of neural activity in the hippocampus after massed or distributed spatial training

**DOI:** 10.1101/2023.05.18.541262

**Authors:** Eleonora Centofante, Luca Fralleoni, Carmen A. Lupascu, Michele Migliore, Arianna Rinaldi, Andrea Mele

## Abstract

Training with long inter-session intervals, termed *distributed training*, has long been known to be superior to training with short intervals, termed *massed training*. In the present study we compared c-Fos expression after massed and distributed training protocols in the Morris water maze to outline possible differences in the learning-induced pattern of neural activation in the dorsalCA1 in the two training conditions.

The results demonstrate that training and time lags between learning opportunities had an impact on the pattern of neuronal activity in the dorsalCA1. Mice trained with the distributed protocol showed sustained neuronal activity in the postero-septal component of the dorsalCA1. In parallel, in trained mice we found more active cells that tended to constitute spatially restricted clusters, whose degree increased with the increase in the time lags between learning trials. Moreover, activated cell assemblies demonstrated increased stability in their spatial organization after distributed as compared to massed training or control condition. Finally, using a machine learning algorithm we found that differences in the number of c-Fos positive cells and their location in the dorsalCA1 could be predictive of the training protocol used. These results suggest that the topographic organization and the spatial location of learning activated cell assemblies might be critical to promote the increased stability of the memory trace induced by distributed training.

## Introduction

Training that includes long intervals between training sessions is termed *spaced* or *distributed training*, and has long been known to be superior to training that includes short intervals, termed *massed training*. Although this phenomenon is ubiquitous over a great variety of organisms and learning tasks ^1–5^ and despite its relevance for education, therapy, and advertising, its neurobiological underpinnings are still poorly understood. Findings in humans as well as in laboratory rodents support the view that *distributed practice* does not affect acquisition of information but it rather yields to better retention of memory at remote time intervals ^6–9^, suggesting that longer time intervals between training stimuli might impact upon the consolidation process by increasing the efficiency of information storage ^10,11^. Experimental evidence supports this view demonstrating that distributed stimuli have a better efficacy in activating molecular pathways important for memory ^10,12^. For example, in *Drosophila* it has been found that distributed but not massed training trials are able to generate distinct waves of MAPK activity ^13^. Similar effects have also been observed after continuous or intermittent 5-HT activation on CREB transcription in Aplysia ^14^.

Individual learning experiences have been shown to induce activity in a sparse population of neurons. This original population of neurons, however, evolves over time and over subsequent stimulations, such changes are crucial for the formation of a stable memory trace ^15,16^. The features of this dynamic process have begun to be outlined. For example it has been shown that the relative excitability of a subset of neurons increases the probability that those neurons will participate to the memory trace ^17^. Interestingly integration at a cellular level between successive stimuli should occur within a limited time window to promote the formation of a more stable memory ^18^. Also relevant to us is the observation that over subsequent trainings, cell ensemble required for retrieval of remote memories differ from those activated by the initial learning stimulus, suggesting a time dependent reorganization of the cellular populations underlying the retrieval of remote memory ^16^. Interestingly, this reorganization process might follow rules promoting topological changes in the cell ensemble embedded into the memory trace to form clusters of cells with coordinated activity ^19,20^.

In this framework we asked whether and how changes in the frequency of training in the spatial version of the Morris water maze (sMWM) might affect cell activity, focusing on the dorsal CA1 (dCA1) of the hippocampus (HPC), with the working hypothesis that different patterns could be observed after massed and distributed learning experiences. We focused on the dCA1 of the HPC for its critical role in spatial memory ^21^ but also because it has been shown to exhibit a high functional heterogeneity ^22–27^. To this aim c-Fos expression in the dCA1 was assessed after learning with the two training protocols, comparing differences in the pattern and stability of the topographical organization of labeled neurons. The analysis demonstrated different regional distribution of c-Fos positive cells in massed and distributed trained mice. Moreover, the findings support the view that distributed training promotes increased clustering and topographical stability in learning activated cells.

## Results

### Massed and distributed trained mice learn to locate the platform successfully

No differences were found in the learning curve of the groups trained with massed (mMWM) or distributed (dMWM) Morris water maze protocol (Fig. 1a) on either latency (two-way repeated measures ANOVA: session F_(5,55)_= 8.322, p< 0.0001; training protocol F_(1,11)_= 0.1476, p= 0.7082; session x training protocol F_(5,55)_= 0.3382, p= 0.8876) or distance (two-way repeated measures ANOVA: session F_(5,55)_= 5.636, p= 0.0003; training protocol F_(1,11)_= 0.02189, p= 0.8851; session x training protocol F_(5,55)_= 0.5393, p= 0.7456), in agreement with what has been previously reported ^9,28^. Further analysis confirmed that both mMWM and dMWM trained mice progressively reduced the latency (one-way repeated measures ANOVA mMWM: F_(5,25)_= 4.194, p= 0.0066; dMWM: F_(5,30)_= 5.706, p= 0.0008) and the distance to reach the platform, (one-way repeated measures ANOVA mMWM: F_(5,25)_= 2.793, p= 0.0388; dMWM: F_(5,30)_= 3.530, p= 0.0125), demonstrating that both groups were able to correctly acquire the task (Fig. 1b,c).

**Figure 1:**
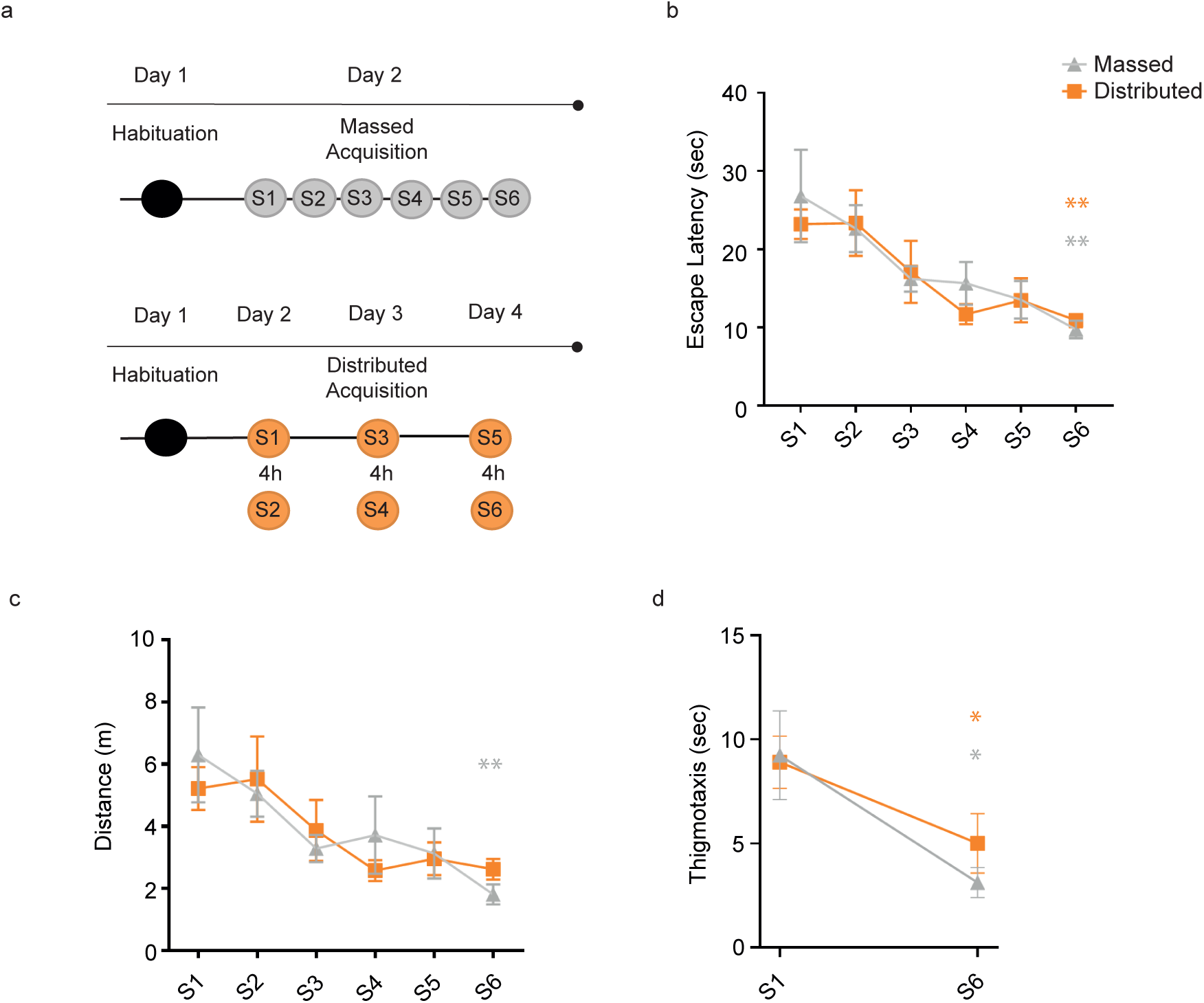
Learning curves for mice trained in the spatial version of the MWM with massed or distributed training protocols. **(a)** Timeline of the two MWM training procedures. **(b)** The graph shows the escape latency to reach the platform during training in the MWM in the massed (grey; N= 6) and in the distributed (orange; N= 7) groups (**p≤ 0.01, S1 vs S6, massed and distributed, Tukey HSD). **(c)** The graph shows the distance travelled during training in the MWM in the massed and distributed groups (**p≤ 0.01, S1 vs S6, massed, Tukey HSD). **(d)** The graph shows the time spent in thigmotaxis in S1 and S6 for both massed and distributed groups (*p≤ 0.05, S1 vs S6, massed and distributed, paired t-test). Graphs represent mean +/− SEM.

A thigmotactic strategy is often deployed by animals in the initial stages of the acquisition as an alternative to a spatial strategy ^29^ therefore the comparison of the thigmotactic behavior in S1 and S6 in the two experimental groups was used to assess increased proficiency at this spatial task. The statistical analysis revealed a significant difference between S1 and S6 for both groups (mMWM: t_(5)_= 2.301, p= 0.0349; dMWM t_(6)_= 2.478, p= 0.0240, paired t-test), demonstrating that with training progression mice reduced the time spent doing thigmotaxis (Fig. 1d).

### Learning induced c-Fos activity in the dCA1 shows regional specificity depending on the training protocol

To investigate changes of c-Fos expression after massed and distributed training in the MWM we focused on the CA1 of the dorsal hippocampus, because it has been shown to be critical for spatial MWM acquisition ^21^. To perform a spatially detailed count of c-Fos positive cells, we draw up to 12 contiguous regions of interest (ROIs) on each section of c-Fos-stained dCA1, as shown in Fig. 2a,b. The mean number of c-Fos positive nuclei per ROI in the dCA1 was higher in trained mice (mMWM: 26.6 ± 5.6; dMWM: 29.3 ± 3.0) compared to controls (20.4 ± 2.4), although the difference did not reach statistical significance (one-way ANOVA: F_(2,24)_ = 1.988, p= 0.158). Next, we analyzed the pattern of expression of c-Fos positive nuclei along the antero-posterior and septo-temporal axes of dCA1, in light of their significant functional heterogeneity ^22–26^ (Fig. 2a,b). To this aim, we generated 3D heatmaps reproducing the number of c-Fos positive cells in each ROI within the dCA1. As shown in Fig. 3a the controls exhibited an overall lower number of activated cells, compared to the massed and the distributed groups, and a fairly homogenous distribution throughout the dCA1. On the contrary, the number of c-Fos positive cells was not evenly distributed in mice trained in the MWM. Moreover, the analysis revealed regional differences depending on the intertrial interval used to train the mice. In particular, the massed trained mice showed higher level of expression in the medio-temporal portion of dCA1 (Fig. 3b) while mice trained with the distributed protocol revealed higher levels in the postero-septal component of the dCA1compared to controls (Fig. 3c). To deepen our analysis and quantitatively compare c-Fos activity in mice trained with the two protocols we grouped the ROIs along the antero-posterior and septo-temporal axes (see methods). The one-way ANOVA revealed significant differences in the mean c-Fos counts in the postero-central and postero-temporal regions (one-way ANOVA for training protocol: F_(2,24)_= 3.15, p= 0.062 for the postero-septal region; F_(2,22)_= 3.618, p= 0.044 for the postero-central; F_(2,22)_= 3.607; p= 0.044 for the postero-temporal; F_(2,24)_= 1.222, p=0.312 for the medio-septal; F_(2,24)_= 1.751, p= 0.195 for the medio-central; F_(2,23)_= 3.216, p= 0.058 for the medio-temporal; F_(2,23)_=0.101; p= 0.905 for the antero-septal; F_(2,23)_= 0.690, p=0.512 for the antero-central) (Fig. 3d). The post-hoc analysis confirmed the significant difference between mice trained with the distributed training protocol in the postero-central and postero-temporal compartments when compared to controls (p<0.01, Tukey HSD) (Fig. 3d). Overall, these findings confirm a functional heterogeneity in the dCA1, demonstrating that the regional pattern of neuronal activity in the dCA1 is sensitive to the time intervals between training sessions.

**Figure 2:**
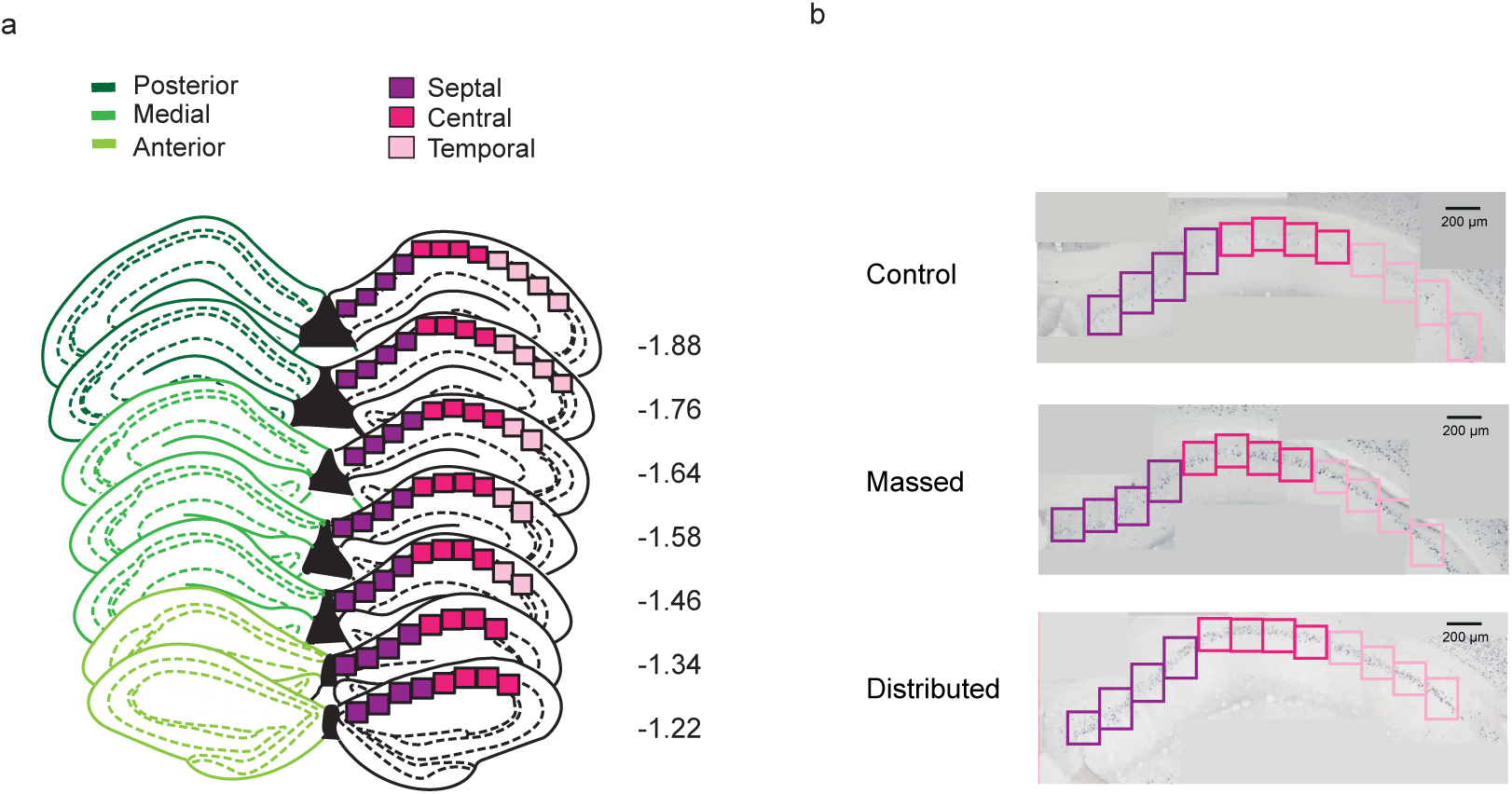
Schematic representation of dCA1 reconstruction. **(a)** Schematic representation of antero-posterior (dark green= posterior; green= medial; light green= anterior) and septo-temporal axes (dark pink= septal; pink= central; light pink= temporal) along dCA1. Numbers indicate anteroposterior coordinates relative to bregma (in mm, according to ^50^. **(b)** Representative posterior dCA1 sections from control, massed and distributed mice, stained for c-Fos. The boxes along the septo-temporal axis represent the position of the ROIs used to count c-Fos positive cells.

**Figure 3:**
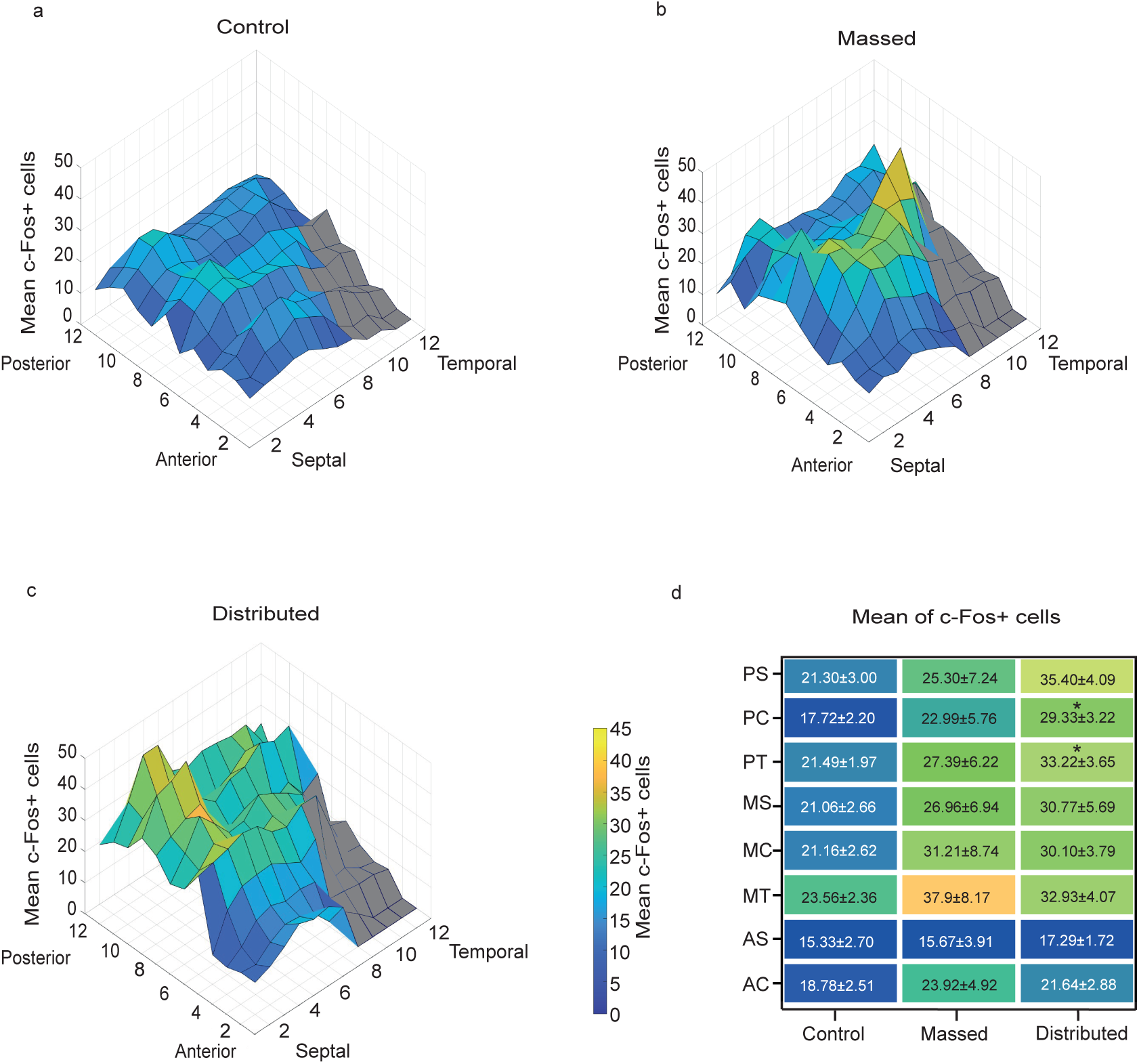
3D representation of c-Fos expression in dCA1 after massed and distributed training. 3D heatmaps representative of the mean of c-Fos positive cells (z axes) along the antero-posterior (12 sections - x axes) and the septo-temporal (8-12 ROIs - y axes) axes in the **(a)** control, **(b)** massed and **(c)** distributed groups. **(d)** Heat maps showing the mean c-Fos positive cell counts in the different subregions (postero-septal (PS), postero-central (PC), postero-temporal (PT); medio-septal (MS), medio-central (MC), medio-temporal (MT), antero-septo (AS), antero-central (AC)) (*p≤ 0.05 distributed vs controls, Tukey HSD). Colored ROIs represent the ROIs used to sample the dCA1.

### Topology of neuronal activity in the dCA1 after massed and distributed training

Next, we verified whether the training protocol could also have affected the topological organization of c-Fos positive cells. As shown in Fig. 4a, the classification of the ROIs based on their c-Fos positive cell number revealed a higher frequency of ROIs in the intervals between 41-80 cells in the distributed compared to controls (*p≤ 0.05; **p≤ 0.01; Mann-Whitney).

**Figure 4:**
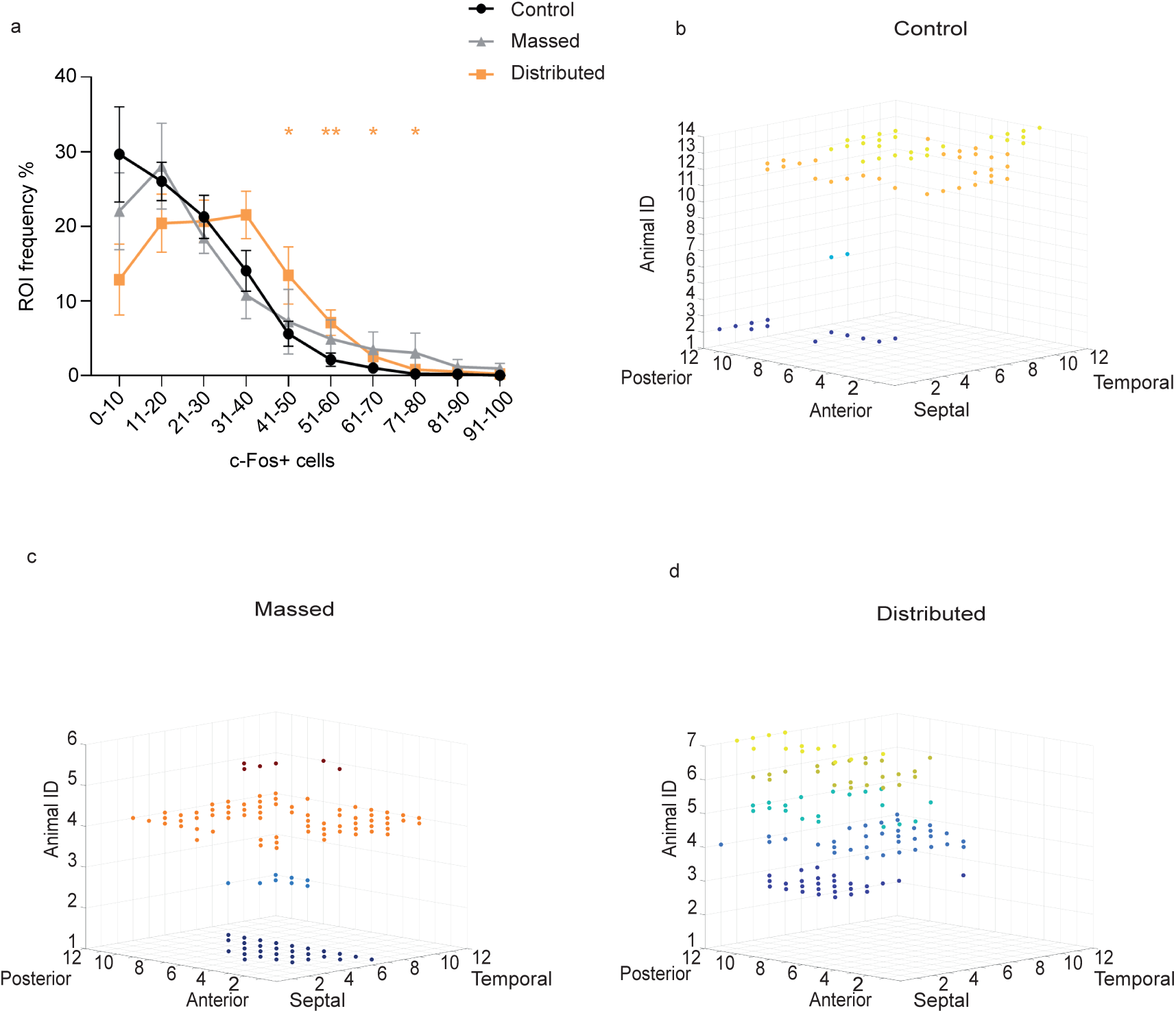
ROIs frequency and 3D distribution of ROIs expressing high number of c-Fos positive cells. **(a)** The graph shows the frequency of ROIs with different number of c-Fos^+^ cells [U=22, (41-50) interval; U=14, (51-60) interval; U=22, (61-70) interval; U=18, (71-80) interval. *p≤ 0.05; **p≤0.01 distributed vs control, Mann-Whitney] **(b-d)** 3D representation of ROIs clusters defined as the ROIs expressing high levels of c-Fos^+^ cells (≧40) neighbored at least 5 similar ROIs (c-Fos^+^≧40) separately for each experimental subject of the **(b)** control, **(c)** massed and **(d)** distributed groups. Each color represents one animal.

Additionally, using a custom made MATLAB script (see methods), we identified, among the ROIs with a high cell count (c-Fos^+^ cells≧40), only those that neighbor (within 5 Euclidean distance) at least five other high-count ROIs, showing that in trained mice these ROIs were in greater numbers (18.83±11.12 massed group; 17.2±4.5 for distributed and 4.9±2.7 for control) (Fig. 4b,c,d), with a significant difference between the distributed and the control groups (t_(19)_= 2.494, p= 0.022 dMWM vs controls; t_(18)_= 1.702, p= 0.106, mMWM vs controls). These results suggest that the distributed training protocol, more efficient in optimizing memory, promotes the clustering of learning-induced active cells within the dCA1.

### Training-induced unique c-Fos expression pattern

Using a custom-made MATLAB script (see methods) we analyzed the activation pattern in the massed and in the distributed groups compared to controls. Considering the mean of c-Fos positive cells per ROI in controls (c-Fos^+^= 23) as a threshold for all the three groups, we searched for ROIs activated over the threshold with the same topological position in at least 5 out of 6 animals in the massed, 6 out of 7 in the distributed and 13 out of 14 in the control group. We could not find any above-threshold ROI sharing the same position in the animals of the control group (Fig. 5a), this held true even considering shared ROIs in only 5 out of 14 mice of the control group. Conversely this analysis revealed that 4 above-threshold ROIs in animals of the massed group and 11 above-threshold ROIs in the distributed group shared the same position (Fig. 5b,c). Coherently with the regional activation induced by the training protocols, these common ROIs were specifically localized in the medial compartment of dCA1 for the massed and in the posterior portion of dCA1 for the distributed group, as shown in the 3D reconstruction in Fig. 5.

**Figure 5:**
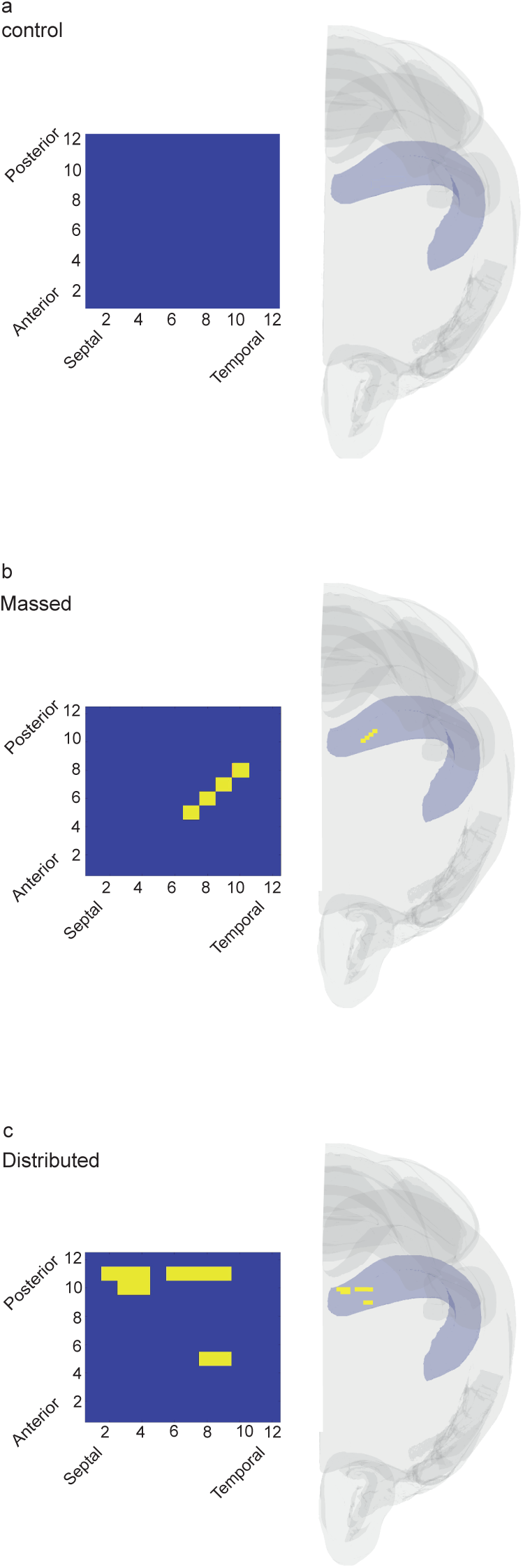
ROIs above threshold with shared position in the different experimental groups. **(a)** Controls, **(b)** massed and **(c)** distributed 3D brains reconstruction showing the localization of the shared above-threshold ROIs (yellow ROIs) within the dCA1.

These results led us to ask whether these characteristic activation patterns could be robust enough to predict the training procedure. We computed accuracy using a Quadratic Discriminant (QD) classifier, a supervised machine-learning algorithm for data classification. When trained with all the variables (see methods), the algorithm showed a remarkable capability to discriminate between massed and distributed trained animals (accuracy= 78%). Notably, when we repeated the procedure adding the control group the level of accuracy decreased (accuracy= 41.4%), pointing out how the absence of a specific c-Fos expression pattern in the control group reduced the discrimination ability of the algorithm. The analysis of feature importance highlighted that the most informative variable was the antero-posterior coordinate (section number) confirming that regional differences in the level of c-Fos activity in the trained animals was a critical feature to distinguish among groups (Fig. 6).

**Figure 6:**
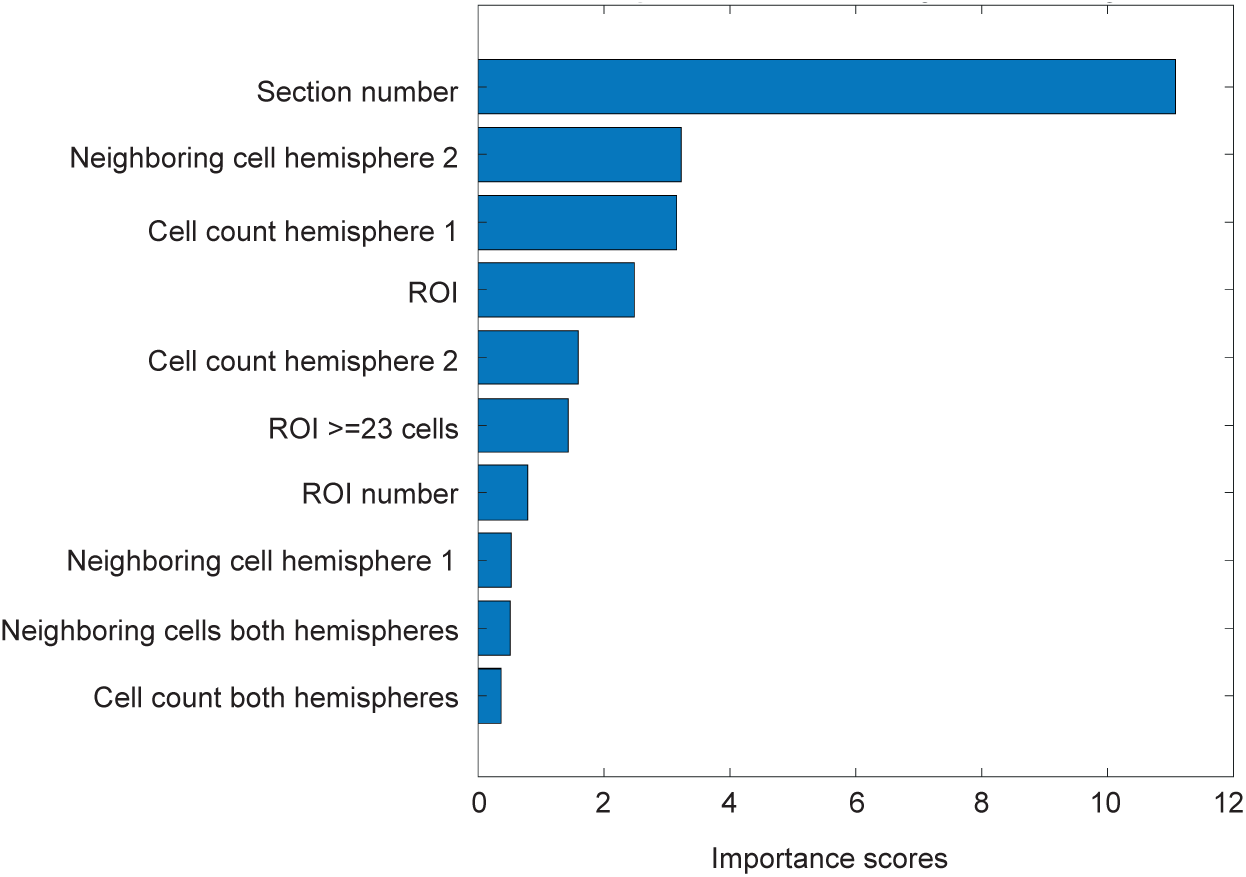
Machine-learning based classification of the training procedure. Histogram depicts the relative importance of the top 10 features (Section number, Neighboring cell hemisphere 2, Cell count hemisphere 1, ROI, Cell count hemisphere 2, ROIs ≥23cells, ROI number, Neighboring cell hemisphere 1, Neighboring cell both hemispheres, cell count both hemispheres) in the Quadratic Discriminant model (Kruskal-Wallis algorithm features importance scores).

## Discussion

In the present study we contrasted c-Fos expression after massed and distributed training protocols in the spatial version of the MWM to outline possible differences in the learning induced pattern of neural activation in the dCA1 of hippocampus in the two training conditions. Three main results were obtained: 1. Distinct regional distribution of c-Fos positive cells after massed and distributed training; 2. Increased clustering of cell activity in mice trained with the distributed as compared to mice trained with the massed protocol; 3. Increased stability across animals of spatially patterned organization of distributed training induced active cells assemblies. Moreover, using a Quadratic Discriminant algorithm we found that differences in the number of c-Fos positive cells and their location in the dCA1 could be predictive of the training protocol used.

It is well known for many types of learning that distributed training, involving longer inter-trial intervals, leads to a more robust memory formation as compared to massed training ^2,5,6,9^. In the present study we compared massed and distributed training in the spatial version of the MWM. Two different training protocols were used, a within day protocol in which the six training sessions are massed with a 5-10 min inter-trial interval, and an across-three-days protocol that has been recently demonstrated to increase memory stability ^9^, reproducing the increased memory performance observed in humans when inter-repetition lags increase ^11^. Independently on the training protocol, mice of the two groups reduced the distance and the time to reach the platform. Consistently with previous evidence ^5,8,9,28,30,31^ distributed training did not affect learning, as there was no difference in terms of the total distance moved and the latency to find the platform as the sessions progressed. These finding are in agreement with the suggestion that the ability of distributed training to increase memory cannot be attributed to an effect of the training protocol on learning, but rather on the consolidation process ^9–11^.

Although we could not find any behavioral differences in the acquisition phase, marked differences could be detected in the pattern of c-Fos expression in the dCA1 of the HPC depending on the training protocol. Moreover the correlation between changes in IEG expression and spatial learning and consolidation is well established ^32–35^.

Analysis of c-Fos labeled cells did not highlight significant differences in the overall number of active cells between the two experimental groups, however we found a change in the regional distribution of c-Fos labeled cells. Our analysis revealed that increasing lags between learning opportunities changes the pattern of activity over the septo-temporal axis as well as over the antero-posterior axis of dCA1. In massed trained mice, in fact, we observed a higher number of c-Fos expressing cells in the medio-temporal component of the CA1, while distributed training induced activity was found mainly in the postero-septal domain of the CA1. Although the hippocampal networks sustaining memory within the CA1 are traditionally viewed as constituted by homogeneous populations of cells, heterogeneity has been demonstrated both at a structural and a functional level. The structural heterogeneity is well established based on the differential projections from the entorhinal cortex, with the lateral part projecting preferentially to the septal component of the CA1 and the medial part projecting mainly to the temporal CA1 (van Groen et al., 2003; Witter et al., 2006). The cellular compactness and width of the deep and superficial CA1 pyramidal cells sublayers also show a regionalization, decreasing and increasing respectively along the septo-temporal axis ^22^. This structural heterogeneity is paralleled by functional differences on the septo-temporal axis. Relevant to us is the finding that cells in the septal component display a higher place field accuracy compared to those in the temporal component ^24^. Such a functional gradient is paralleled by an opposite radial gradient in the place field stability ^23^ and in the recruitment of CA1 pyramidal neurons during HPC sharp wave ripples ^38^, suggesting a higher dynamicity and accuracy of cells in the temporal CA1 as compared to those in the septal component that would show a coarse representation of the environment but with a higher steadiness ^22^. Interestingly the different pattern of activity in mice trained with the massed or the distributed protocols suggest a progressive shift along the septo-temporal axis with the increased time lag between learning opportunities. The temporal component is indeed activated after massed training, whereases the septal is activated after a distributed training (Fig. 3). This result would be consistent with behavioral evidence in rodents and humans that memory representation after distributed learning is more stable than after massed learning ^9,11^.

The second interesting observation raised by the quantification of c-Fos labeling was that the topographic distribution of learning activated cells in the dCA1 varies depending on the training protocols. In particular the analysis performed revealed a group difference in the number of ROIs expressing higher c-Fos labeling. This observation, together with the clustering of ROIs expressing a high number of c-Fos positive cells in the distributed groups, as compared to control mice or mice trained with the massed protocol, strongly suggests a training protocol dependent clustering of learning activated cells. Two different lines of evidence support the possibility of a learning induced clustering of cell activity. A handful of electrophysiological studies for example reported a cluster type organization of place cells, demonstrating that neighboring pyramidal cells in the CA1 have overlapping place fields (Eichenbaum et al., 1989; Deadwyler et al., 1996; Hampson et al., 1999), but see also ^42^. More relevant are findings relative to the organization of IEGs labeled cells after training, demonstrating increased IEGs activity in adjacent cells, in the CA1, after exposure to a novel environment and tighter clusters when exposure to the same environment is repeated ^43,44^. A similar learning dependent increase in clustering of active cells was observed also in the dorsal striatum as instrumental learning progresses ^20^. Although a causal role of such clustering in memory formation has never been established, our findings support this hypothesis, by showing that it is more pronounced increasing the time interval between sessions when the stability of the memory trace is enhanced.

Finally, we also found a difference in the stability of the topographic distribution of activated ROIs in the three experimental groups. A few studies addressed the question relative to the topography of learning activated cells in the HPC as well as in other brain regions ^45–47^. The increased topographic stability of active cells assemblies we found comparing the two training protocols is in agreement with previous evidence demonstrating a spatially patterned organization of amygdala neurons activated by training in a fear conditioning task ^45,46^ as well as fMRI data reporting a stable topography of activation for specific memories in the human HPC ^48,49^. Thus our data support the hypothesis that increased stability of the memory trace might depend on the spatial location of active cell assemblies within dCA1 ^45,49^. Finally, we were able to discriminate mice trained with the different training protocols based on a quadratic discriminant algorithm for data classification, using information relative to the pattern of expression of learning induced c-Fos positive cells. These findings suggest that optimization of memory might depend upon specific hippocampal neuronal population codes that vary depending on the stability of the memory trace.

It has been hypothesized that the memory stability induced by distributed training depends on a more efficient stabilization of the memory trace (Smolen et al., 2016). The evidence presented in this study suggests that the topographic organization and the spatial location of learning activated cell assemblies might be relevant to this process. Moreover, it supports the view that activity in the septal component of the dCA1 might be important in determining the stability of the representation.

## Materials and Methods

### Subjects

The experiments were conducted on naïve CD1 male mice (Charles River, Italy). Mice were 8 weeks old, weighting 34-38 g, at the onset of the experiments. Animals were always housed in groups of three mice in standard cages (26.8 × 21.5 × 14.1 cm), with water and food ad libitum, under a 12 h light/dark cycle (7am/7pm) and constant temperature (22 ± 1 °C). Behavioral training and testing were conducted during the light period (from 9:00 am to 5:00 pm). The maximum effort was made to minimize animal suffering. Procedures were conducted under the authorization N. 450/2018-PR from Italian Ministry of Health, according to Italian (DL.116/92) and European laws and regulations on the use of animals in research, and NIH guidelines on animal care.

### Morris water maze

The apparatus consisted of a circular pool (110 cm diameter and 40 cm high) filled up to 5 cm from the edge with water at 22 +/− 1°C. To allow mice tracking, water was black-colored with non-toxic paint (Giotto, Italy). Black curtains surrounded the pool, and several visual cues were attached on them at the distance of round 50 cm from the pool. A white light (100W) was positioned above the apparatus facing the ceiling. Four additional lights (1.9 W) were positioned facing the cues to light them up. The pool was ideally divided into four identical quadrants. Four starting positions (labeled N, S, E, and W) were located equidistantly around the edge of the maze. A Plexiglas platform (10 cm diameter), covered with wire mesh to avoid slipping, was positioned 20 cm away from the pool wall in one of the four quadrants. The quadrant where the platform was kept during the training was randomized and was named the “Target” quadrant, whereas the other three quadrants were named “Right”, “Left” and “Opposite”, relative to the Target.

Mice of each group were handled for 3-4 min/day, for 5 consecutive days, before the beginning of the experiments. The spatial version of the Morris Water Maze task (MWM) consisted of two different phases: familiarization and training (Fig. 1a). The familiarization phase was the same for both the massed (mMWM) and the distributed (dMWM) protocols and consisted of one session of three consecutive trials (intertrial interval: 20 s). During the familiarization phase, no cues were attached on the curtains and the platform protruded 1.5 cm above the water surface. During the training phase, the platform was submerged 0.5 cm beneath the surface of the water and visual cues were attached on the black curtains surrounding the pool. The mMWM consisted of six consecutive training sessions (intersession interval: 10-15 min) of three trials (intertrial interval: 20s), whereas dMWM consisted of six training sessions distributed over 3 days, with 2 sessions per day (intersession interval: 4 h) of three trials (intertrial interval: 20 s) ^9^. In each trial, animals were randomly released from one of the three non-target quadrants. Mice in mMWM were kept in their holding cage for 1h before the starting of the training sessions, whereas mice in dMWM were kept in their holding cage for 1h before each training session. During intersession intervals, mMWM mice returned to their holding cage until the next session. Control animals were kept in their holding cage for the whole length of training sessions.

All trials were recorded by a camera located over the pool and videos were acquired and analyzed by an automated tracking system (AnyMaze 5.0, Stoelting). Escape latency (in seconds) and distance travelled to reach the platform (in meters) were scored. Thigmotaxis was defined as the swimming time within the zone closer to the pool wall (10 cm from pool wall).

### Immunohistochemistry: Fos staining

One hour after completing the last training session in the MWM, each animal was deeply anaesthetized with a mixture of Zoletil (500 mg/kg, Virbac Italia) and Xylazine (100 mg/kg, Bayer, Germany) and transcardially perfused with 40 ml of saline solution (NaCl 0.9%) followed by 40 ml of 4% formaldehyde in PBS (4°C). Brains were rapidly removed and post-fixed for 24 h in 4% formaldehyde and then transferred to 30% sucrose in PBS until sectioning. 40 μm-coronal sections were obtained using a cryostat (Leica CM 1520, Leica Microsystems, Wetzlar, Germany) and were stored at −20 °C in cryoprotectant solution. For the c-Fos-immunoreactivity (c-Fos-IR), free-floating sections were incubated for 5 min in 3% hydrogen peroxide in PBS and rinsed three times in PBS containing 0.1% Triton X-100 (PBST). After 1 h of incubation in PBST containing 1% BSA and 1% NGS sections were incubated overnight in anti-phospho-c-Fos rabbit monoclonal antibody (5348S; Cell signaling Technology, USA) diluted 1:8000 in PBST– BSA–NGS, at 4 °C and with constant orbital rotation. Sections were washed three times in PBST and incubated in biotinylated secondary antibody diluted 1:500 in PBST-1% BSA (goat anti-rabbit IgG; Vector Laboratories, USA) for 2 h at room temperature. After three washes in PBST, sections were transferred for 1 h in avidin–biotinylated peroxidase complex diluted 1:500 in PBST (ABC Kit; Vector Laboratories, USA) and rinsed three times in PBS. Finally, the reaction was visualized using nickel intensified diaminobenzidine (DAB peroxidase substrate kit, Vector Laboratories, USA). The reaction was stopped after exactly 4 min by washing with 0.1 M PBS (pH 7.6). Sections were mounted on slides, dehydrated through a graded series of alcohols, cleared and coverslipped. Sections from groups to be directly compared were processed at the same time and using the same conditions and reagents to reduce variability. In all experiments, the number of cells displaying c-Fos immunoreactivity was measured in dCA1, taking twelve alternating brain sections (from −1.22mm to −1.92mm relative to bregma), encompassing the whole antero-posterior axis. Regions were defined according to the mouse brain atlas ^50^.

### Digital reconstruction and c-Fos quantification

Digital images were acquired at 10X magnification, using a microscope (Nikon Eclipse 80I) equipped with a CCD camera. After images acquisition, single images of 2560 × 1920 pixel were stitched together in a mosaic view of the entire dCA1 region and counting of the stained nuclei was carried out using the public domain software ImageJ (http://rsb.info.nih.gov/ij/). Briefly, for each section, stained nuclei were automatically detected based on their intensity of staining relative to background and their size. Detailed counting of c-Fos positive cells was performed drawing from 8 to 12 contiguous regions of interested (ROIs), covering the whole dCA1 on each section. Each ROI had a 200 *µm* width and ROIs were placed consecutively along the dCA1 septo-temporal axis. The anchor point for the first ROI was always set at the beginning of the dCA1 along the midline of the brain, using the ventricle as the starting point marker. The dCA1 was covered by 8 ROIs in the most anterior sections, 10 ROIs in the medial region, whereas in the most posterior region was covered by 12 ROIs per section. Although missing ROIs were present in every animal the mean number of ROIs sampled was similar in the different groups (control: 87±5.7; massed: 89.8±10.8; distributed: 90.4±7.4). ROIs in the whole dCA1 were also classified based on their number of c-Fos positive cells, according to the following intervals: 0-10; 11-20; 21-30; 31-40; 41-50; 51-60; 61-70; 71-80; 81-90; 91-100 c-Fos+ cells per ROI. For each experimental subject, the frequency of ROIs in each interval was then calculated and reported as percentage of the total number of ROIs for that subject.

Regional c-Fos expression was evaluated dividing the whole dCA1 in eight different subregions: antero-septal (AS), antero-central (AC), medio-septal (MS), medio-central (MC), medio-temporal (MT), postero-septal (PS), postero-central (PC), postero-temporal (PT). Each subregion includes 16 ROIs except for the medio-temporal compartment (10 ROIs), according to the dCA1 shape.

### Analysis of c-Fos expression pattern

Data obtained from digital image reconstruction and c-Fos quantification were exported in a tab-delimited spreadsheet and imported in MATLAB. MATLAB heatmap representation was used to obtain the 3D heatmap representation of c-Fos positive cells for each ROI and section (Fig. 3). Custom-made scripts (Mata and Matrixa) were used to identify the ROIs with a cell count equal or above 40 cells (threshold chosen based on the differences observed in the ROIs frequency analysis) and neighboring other ROIs (within 5 Euclidean distance) with the same cell count (Fig. 4 b,c,d). Finally custom-made scripts (Histcut_23; Pixels_m_23; Common_23) were used to highlight the ROIs with a cell count equal or above 23 (mean of c-Fos positive cells per ROI in control animals) and with a shared position within each experimental group (Fig. 5). All scripts are included as supplementary materials.

### Machine learning based classification

A MATLAB based library was used to train the classification model (https://www.mathworks.com/help/stats/choose-a-classifier.html). The model is a supervised machine learning algorithm called Quadratic Discriminant (QD) that is based on discriminant analysis learning.

We applied the QD classifier adopting the function cross_val_score with cv = 10 to perform 10-fold cross-validation (available in MATLAB R2022a). The classifier was trained using the information from each subject about the c-Fos positive cells number along the anterior-posterior axis (section number), c-Fos positive cells number along the septo-temporal axis (ROI number), total number of c-Fos positive cells from each hemisphere (cell count hemisphere 1; cell count hemisphere 2), total number of c-Fos positive cells averaged between the two hemispheres (cell count both hemispheres), the presence and the number of the ROIs above threshold (ROIs expressing c-Fos^+^ cells≧ 23), number of the ROIs above threshold adjacent to each other from hemisphere 1, 2 and the average between the two hemispheres (neighboring cell hemisphere 1; neighboring cell hemisphere 2; neighboring cell both hemispheres). To calculate feature importance in classifier decision, we used the attribute feature selection that expresses the fraction of relative importance for each feature.

### Code availability

Codes for these experiments are available in the supplementary material.

### Data collection and statistical analysis

All data are presented as mean ± standard error. For the MWM experiment, escape latency and distance travelled were analyzed using both two-way and one-way repeated measures ANOVA with session (six levels: session 1 to 6) as repeated measure. Time spent in thigmotaxis between S1 and S6 was analyzed by using a two-tailed paired t-test. A one-way ANOVA was used to compare c-Fos expression along the septo-temporal and the antero-posterior axes in mice from the massed and distributed trained groups, followed by Tukey HSD multiple comparison test when appropriate. ROIs frequency intervals were analyzed using a nonparametric analysis (Mann-Whitney U test).

## Supporting information

Matlab codes

## Acknowledgements

This research was supported by a NARSAD independent investigator grant from the Brain and Behavioral Research Foundation (to A.M.), the AARG-NTF-22-971925 grant from Alzheimer’s Association (to A.R.), a HBP (to A.M. and A.R.) and from the European Union’s Horizon 2020 Framework Programme for Research and Innovation under the Specific Grant Agreement numbers 945539 (Human Brain Project SGA3) (to EC, MM, and CAL).

## Authors contributions

All authors contributed to the study conception and design. Material preparation, data collection and analysis were performed by E.C., L.F., and C.A.L. The first draft of the manuscript was written by E.C., A.R. and A.M. all authors commented on previous versions of the manuscript. All authors read and approved the final manuscript.

## Data availability statement

All data supporting described findings can be obtained from the corresponding author (A.M.) upon reasonable request.

