## Supplementary material for "Specific patterns of neural activity in the hippocampus after massed or distributed spatial training": Matlab codes

**Mata**

```
[r,c]=find(a>40);
mata=zeros(size(a);
for i=1:size(a,1)
    for j=1:size(a,2)
        dist=[];
        if a(i,j)>40
            mata(i,j)=2
            for k=1:size(r,1)
                if sqrt((r(k)-i)^2+(c(k)-j)^2)>0
                    dist=[dist sqrt((r(k)-i)^2+(c(k)-j)^2)];
                end
            end
            i
            j
            a(i,j)
            dist
            size(find(dist<=5),2)
            if size(find(dist<=5),2)>=5
                mata(i,j)=size(find(dist<=5),2)
            end
        end
    end
end
imagesc(mata)
```

**Matrixa**

```
function [mata,nra]=matrixa(a)
[r,c]=find(a>60);
nra=0;
mata=zeros(size(a));
for i=1:size(a,1)
    for j=1:size(a,2)
        dist=[];
        if a(i,j)>40
            mata(i,j)=2;
            for k=1:size(r,1)
                if sqrt((r(k)-i)^2+(c(k)-j)^2)>0
                    dist=[dist sqrt((r(k)-i)^2+(c(k)-j)^2)];
                end
            end
            %i
            %j
            %a(i,j)
            %dist
            %size(find(dist<=5),2)
            if size(find(dist<=5),2)>=3
                mata(i,j)=size(find(dist<=5),2);
```

```
                nra=nra+1;
            end
        end
    end
end
end
```

**Histcut_23.m**

```
B=B_X_m;
[counts2 binCenters2] = hist(B(B(:)~=0), 20);
counts3=counts2/sum(counts2);
i=1;
while i<=size(binCenters2,2)
    s=sum(counts3(1:i));
    if s>=0.75
        break;
    end
    i=i+1;
end
figure;imagesc(B,[0 121]);
hold on;
coordzone_B_X_m_23=[];
for k=1:size(B,1)
    for j=1:size(B,2)
        if B(k,j)>=23
```

**Pixels_m_23.m**

```
MATPIXSP=zeros(size(B_X_1));
MATPIXM=zeros(size(B_X_1));
for i=1:12
    for j=1:12
        [tf, index]=ismember([i j],coordzone_B_X_m_23,'rows');
        if (tf==1)
            MATPIXSP(j,i)=MATPIXSP(j,i)+1;
        end
        if not(isequal(B_X_m(j,i),0))
            MATPIXM(j,i)=MATPIXM(j,i)+1;
        End
figure;imagesc(MATPIXSPT)
figure;imagesc(MATPIXM)
hold on;
for k=1:12
    for j=1:12
           a=0;
        if B_X_m(k,j)>=23
                a=a+1;
           end
           %if MATPIXM(k,j)==a && a~=0
           if MATPIXM(k,j)==a && a>=3
            plot(j, k, 'r+', 'MarkerSize', 5, 'LineWidth', 3);
        end
```

```
    end
end
MATPIXSPT2=zeros(size(B_1_1));
for k=1:12
    for j=1:12
          a=0;
        if B_X_m(k,j)>=23
               a=a+1;
          end
if MATPIXM(k,j)==a && a>=6 (spaced) a>=5 (massed) a>=13; 5; 6(control)
           MATPIXSPT2(k,j)=a;
        end
    end
end
figure;imagesc(MATPIXSPT2)
```

**Common_23**

```
MATPIXSP10=zeros(size(B_X_1));
MATPIXM=zeros(size(B_X_1));
for i=1:12
    for j=1:12
        [tf, index]=ismember([i j],coordzone_B_X_m_23,'rows');
        if (tf==1)
           MATPIXSPX(j,i)=MATPIXSPX(j,i)+1;
          if MATPIXSPT(j,i)==1
           MATPIXSPX(j,i)=2;
        end
        end
    end
end
```
